# Simulated poaching affects global connectivity and efficiency in social networks of African savanna elephants - an exemplar of how human disturbance impacts group-living species

**DOI:** 10.1101/2020.08.24.252536

**Authors:** Maggie Wiśniewska, Ivan Puga-Gonzalez, Phyllis Lee, Cynthia J. Moss, Gareth Russell, Simon Garnier, Cédric Sueur

## Abstract

Selective harvest, such as poaching, impacts group-living animals directly through mortality of individuals with desirable traits, and indirectly by altering the structure of their social networks. Understanding the relationship between the structural network changes and group performance in wild animals remains an outstanding problem. To address this knowledge gap, we evaluate the immediate effect of disturbance on group sociality in African savanna elephants — an example, group-living species threatened by poaching. Drawing on static association data from one free ranging population, we constructed 100 virtual networks; performed a series of experiments ‘poaching’ the oldest, socially central or random individuals; and quantified the immediate change in the theoretical indices of network connectivity and efficiency of social diffusion. Although the virtual networks never broke down, targeted elimination of the socially central conspecifics, regardless of age, decreased network connectivity and efficiency. These findings hint at the need to further study resilience by modeling network reorganization and interaction-mediated socioecological learning, empirical data permitting. Our work is unique in quantifying connectivity together with global efficiency in multiple virtual networks that represent the sociodemographic diversity of elephant populations likely found in the wild. The basic design of our simulation platform makes it adaptable for hypothesis testing about the consequences of anthropogenic disturbance or lethal management on social interactions in a variety of group-living species with limited, real-world data.

**Author Summary:** We consider the immediate response of animal groups to human disturbance by using the African savanna elephant as an example of a group-living species threatened by poaching. Previous research in one elephant population showed that poaching-induced mortality reduced social interaction among distantly related elephants, but not among close kin. Whether this type of resilience indicates that affected populations function similarity before and after poaching is an open problem. Understanding it is important because poaching often targets the largest and most socially and ecologically experienced group members. Drawing on empirical association data, we simulated poaching in 100 virtual elephant populations and eliminated the most senior or sociable members. Targeted poaching of sociable conspecifics was more impactful. Although it did not lead to population breakdown, it hampered theoretical features of interspecific associations that in other systems have been associated with group cohesion and the efficiency of transferring social information. Our findings suggest that further inquiry into the relationship between resilience to poaching and group performance is warranted. In addition, our simulation platform offers a generalizable basis for hypothesis testing in other social species, wild or captive, subject to exploitation by humans.

## Introduction

In group-living animals, from insects to mammals (1, 2), interactions among conspecifics with diverse social roles (3–5) impact individual survival (6–9), reproductive success (10–12) and adaptive behaviors (13–16). In species with complex organization characterized by flexible aggregates of stable social units (17–19), the loss of influential group members through natural or anthropogenic causes can be detrimental to surviving conspecifics (20–22) and to entire populations (23, 24). Unlike natural phenomena, such as fire (25, 26), harvest is intrinsically nonrandom (27–29). For instance, poachers profiting from pet trade prefer to capture immature individuals as the most desirable commodity (30), eliminating gregarious ‘brokers’ of social interactions (31, 32). As another example, trophy hunters target individuals with prominent features, such as elephants with big tusks (33, 34), killing the oldest and socioecologically experienced conspecifics (35–37).

Animal social network analysis (ASNA) can be a powerful tool in demonstrating how selective elimination of individuals with key social roles impacts closely knit animal groups. Quantifying relationships between members of a group as ‘networks of nonrandomly linked nodes’ (38, 39) has revealed that while some disturbed groups break down (40, 41), others stay connected (20, 42). Understanding whether the relationships in remaining groups operate as prior to disturbance is based on a small number of studies. In an instance of captive zebra finches, group foraging ability decreased following repeated social disturbance (43). In simulated primate groups, network disturbance led to a decrease in its global connectivity and the efficiency of social diffusion but did not lead to group fragmentation (44). These indices depend on network structure; are based on an assumption that transmissible currency, such as information, diffuses through network links (45); and have been related to cohesion, the transfer of social currency and robustness to loss of influential conspecifics in animal groups (46–48). In light of the anthropogenic impact on ecological communities (49–52), evaluating the relationship between post-disturbance social structure and limitations to social resilience vis-à-vis group performance in natural animal systems is becoming increasingly important (20, 53).

To explore this relationship, we considered the African savanna elephant (*Loxodonta africana*) — a group-living species threatened by poaching (54–56). Elephant social organization consists of several tiers, ranging from transitional clans and bonded groups of distant kin, to matrilinear core units of adults and their immature offspring (57); or flexible groups of post-dispersal males of varying ages and kinship (36). While immature elephants frequently engage in affiliative interactions (58, 59), mature individuals are more experienced about resource distribution and phenology (60, 61) and about social dynamics (62–64). The interactions among individuals with diverse social roles across social tiers manifests as fission-fusion dynamics in response to changing sociophysical landscape (19, 65). Poaching (which during the militarized wave of the past decade eliminated large subsets of populations including mature and immature elephants (66)), impacts demography (67), resource acquisition (68, 69) population genetics (70) and various social behaviors (71, 72) in affected populations.

Evidence from ASNA of data spanning periods of low and high poaching in one free-ranging population revealed that the composition and association patterns within matrilines were conserved among close but not distant surviving kin. This outcome suggests clan-level impact of poaching on network structure and resilience, with little detrimental effect at the bonded group-or core unit-levels (73). Whether changes in network structure in elephants relate to group functionality is difficult to test directly. However, quantifying network connectivity together with global efficiency while simulating poaching may shed new light on the theoretical capacity for dissemination of social currency and the limitation to social resilience in disturbed populations. These insights may eventually inform our understanding about the mechanisms of group performance, as well as the efforts to mitigate human-elephant conflict (74, 75) and conserve this economically important but endangered, keystone species (76, 77).

We characterized the immediate effect of eliminating the most influential individuals on the global structure of simulated, social networks. We used a static set of empirical association data on one free-ranging elephant population from Amboseli National Park (NP) in Kenya (78) because continuous data featuring network reorganization after poaching, necessary to parametrize time-varying models, do not yet exist for wild elephants. Initially, we assembled one social network using an Amboseli dataset and conducted a series of ‘poaching’ experiments by either incrementally removing 1) the oldest elephants as presumably the most experienced and prone to poaching, or topologically central individuals as the most sociable network members (79, 80); or 2) by removing individuals randomly (41, 81). To quantify network-wide structural changes, we evaluated four theoretical indices expressing network-wide connectivity (i.e., clustering coefficient and modularity, dependent on local neighborliness or global partitioning, respectively); as well as the efficiency of social diffusion (i.e, diameter and global efficiency, based on the distance or pervasiveness of diffusion, respectively) (47). To set these results in the context of a large-scale variation in demography and social interactions found in real elephant populations, we generated 100 distinct, virtual populations modeled on demographic trends in empirical data. To simulate social network formation in these populations, we built a spatiotemporally nonexplicit, individual-based model with rules informed by empirical associations (57, 78). The steps of assigning social influence, conducting deletion experiments and quantifying deletion effects were as mentioned earlier.

We hypothesized that elimination of the most influential individuals, defined according to their age category or network position would lead to a decrease in global network connectedness and efficiency. Specifically, we predicted that relative to random deletions, targeted removal of the most central or mature individuals would result in a decrease in global clustering coefficient and efficiency, and an increase in diameter and modularity. We also anticipated a worsening in these outcomes as a function of the proportion of deleted individuals, resulting in an eventual network breakdown. This set of findings would be an indication of increased subgrouping at the population level, fewer interactions with immediate social partners and fewer pathways for timely and fault-tolerant transfer of social currency.

Partly consistent with our expectations, our work shows that targeted ‘poaching’ of the most sociable elephants was more impactful on simulated network structure than elimination of senior network members. Although it was not parameterized to reflect the rate of ‘poaching’ events in absolute time, and cannot be used to inform response to poaching after network reorganization, our work offers a novel perspective on the immediate response to disturbance in a large number of sociodemographically diverse populations with experience of poaching-like stress. Keeping in mind the delimitations of our approach, we interpret our findings in the context of a common behavioral repertoire in wild elephant populations and offer insights about how our findings may potentially help view natural populations subject to poaching. Finally, we consider the utility of our simulation platform as a generalizable tool for testing hypotheses about the disturbance of social dynamics in other species that facilitate ecosystem functioning or impact human welfare (82, 83).

## Materials and Methods

We performed a series of deletions using one social network derived from association data on a free-ranging elephant population and 100 virtual networks mimicking the empirical one. Details of these experiments and underlying assumptions are described in the following subsections.

### Empirical data

#### Specifying empirical population composition

To gather baseline information about demography and social interactions characterizing elephant sociality, we considered two dyadic association datasets from Amboseli NP originally published elsewhere (78). We assume that these datasets, collected at vantage points where different social units converge to drink, capture of a range of social processes including events that required group cohesion and transfer of information (e.g., conflict avoidance in a multigroup gathering at a waterhole requires learning and recall about which conspecifics to affiliate with and whom to avoid (84)).

During the original data collection, the authors inferred proximity-based, dyadic associations at two social tiers: among individuals within 10 separate, core groups (within core group -WCG) and between 64 core groups (between core group - BCG), where each group was treated as a single social entity. However, we had a different goal — to examine population-wide dynamics. To represent associations that occurred within each core group in the population, we used the unaltered WCG association data according to the following association index (AI) formula: *AI_i,j_ = x_i,j_ / (x_ij_ + d + (n - d - x_i,j_))*. In this formula, *x_ij_* is the number of times individuals *i* and *j* were seen together; *d* is the number of times neither individual was seen; *n* is the total number of times a group was observed; and by extrapolation (n *-d - x_ij_)*) represents the number of times either individua*l i* or *j* was seen. To express the interactions occurring between individuals from different core groups, we assembled a dyadic association matrix by combining the WCG and BCG data as (85) (85).

Although the original dataset included 64 groups, we could only focus on 10 groups for which both WCG and BCG data were available (labeled AA, CB, DB, EA, EB, FB, JAYA, GB, OA, and PC). To reflect the typical, multi-tier structure of an elephant society (57), we aggregated the 10 core groups into eight bond groups [i.e., B1 (core group AA, including 10 individuals); B2 (FB, 6); B3 (EA, 9 and EB, 10); B4 (DB, 4); B5 (CB, 6 and OA, 10); B6 (GB, 11); B7 (PC, 9); and B8 (JAYA, 8)] and three clan groups [i.e., K1 (bond groups B1, B2, B3 and B4); K2 (B5, B6 and B7); and K3 (B8)] using genetically determined relatedness indices and long-term, behavioral associations inferred by the authors (78).

#### Inferring population-wide social interactions and assembling one social network based on empirical association data

We calculated the fraction of all sightings when an individual *i* from core group *G* was seen in that group according to the following formula: *f_i,G_ = average n_i,j,G_ / (n_G_ – average d_i,j,G_)* where the averages are over all the other individuals *j* in group *G*. In this formula, *n_i,j,G_* represents the number of times individuals *i* and *j* were seen within group *G*; *d_i,j,G_* is the number of times neither individual *i* nor individual *j* was seen within group *G*; and *n_G_* is the number of times group *G* was observed. The denominator is, therefore, the average number of times group *G* was observed with either individual *i*, individual *j* or both present; and *f_i,G_*, which falls in the interval {0,1}, can be thought of as the average fraction of these occasions when they were both present or an index of the overall sociability of individual *i*. This process was repeated for every individual in the population.

Using the information available for the BCG association data, we calculated the fraction of all sightings when group *G* was seen with group *B* according to the following formula: *f_G,B_* = *n_G,B_* / (*n_G_* + *n_B_* – *n_G,B_*). Here, *n_G,B_* indicates the number of times groups *G* and *B* were seen together; *n_G_* indicates the number of times group *G* was seen without group B; and *n_B_* indicates the number of times group *B* was seen without group G. Thus the denominator is the number of times groups *G* and *B* were seen individually. This process was repeated for every pair of groups in the population and can be thought of as the probability of seeing a given pair of groups together. We then derived a symmetric, weighted matrix consisting of probabilities of dyadic associations between individuals from two different groups, for instance, individuals *i_G_* and *aB* from groups *G* and *B* respectively, by using the following formula: *p*(*iG*, *a_B_*) = *f_i,G_* × *f_a,B_* × *f_G,B_*. Finally, using this matrix, we constructed a population-wide network of associations or links.

#### Quantifying social influence in empirically based social network

To identify influential network members serving as social centers and intermediaries (86), we quantified each individual’s betweenness and degree scores (80). Given that these metrics were highly correlated, we used betweenness going forward as particularly suitable for questions about global connectivity and more importantly the efficiency of social diffusion in a society with fission-fusion dynamics (48, 87). From this point onward we often refer to individuals with high betweenness scores as the most central individual. To include age as a form of social influence due to presumed disparity in socioecological experience between mature versus immature individuals, we considered four age categories. They included young adults, prime adults, mature adults and the matriarchs (88). Betweenness and age category were not correlated.

Their definitions are detailed in Table 1.

**Table 1.** Definitions of social influence metrics and network indices used in this publication, as well as expected outcomes for weighted (W) and unweighted indices measured after incremental deletion of the most socially influential individuals in targeted deletions, or in random deletions without consideration for their social influence.

| Individual level metric | Definition |  |
| --- | --- | --- |
| Betweenness | The number of shortest paths<sup>1</sup> passing through an individual. High value indicates high social interconnectedness and thus important theoretical role that an individual has in the exchange of social currency, such as information (97,98) |  |
| Age category | A segment of the population within a specified range of ages, including: 1) young adults (individuals <12 and < 20 years old); 2) prime adults (20-35); 3) mature adults (>35); 4) the matriarchs (the oldest or most dominant females in the core group)) used when categorical consideration of age is desired, or when data on absolute age are not available; in the empirically based population the age ranges were based on year of birth; in the virtual populations, the age range distribution was modeled to parallel the empirical distribution of ages (78,88) |  |
| Network level index | Definition | Predictions |
| Clustering coefficient | The number of triplets (where any set of three individuals are connected by either two or three links, referred to respectively as open and closed triplets, respectively) divided by the total possible number of triplets. High values have been associated with high group cohesion, little subgrouping, and resilience against disturbance-induced breakdown (39,80,99) | deletion proportion:<br>0 > 0.2<br>deletion type:<br>random > targeted |
| Diameter W | The path with the maximum weight <sup>2</sup> among the shortest path lengths across all dyads. High values have been associated with low degree of cohesion potentially impeding rapid transmission of information (39,41,80) | 0 < 0.2<br>random < targeted |
| Global efficiency W | The inverse of the network's global efficiency: which measures the ratio between the total number of individuals and links multiplied by the network diameter <sup>3</sup>. High values have been associated with high probability of social diffusion in a group and thus important theoretical role in efficient transmission of information (95,100). | 0 > 0.2<br>random > targeted |
| Modularity W | The density of links within a module in a weighted network relative to the density of links between modules. High value indicates low group cohesion with cohesive subgroups, and susceptibility to breakdown after disturbance (101–103). | $0 < 0.2$<br>random < targeted |

<sup>1</sup>Shortest path - the path with the minimum number of links between any pair of individuals (95,96). <sup>2</sup>Path weight - the inverse of the weight of a link, where links with highest weights are equivalent to shortest paths (48). <sup>3</sup>Diameter - the longest among the shortest path lengths in a network (39).

#### Conducting deletions using empirically based social network

To assess how disturbance affects global structure in elephant social networks and determine the level of stress that would bring about network fragmentation, we carried out a sequence of targeted deletions by selecting 20 percent of the oldest or most central network members (together referred to as ‘*deletion metrics*’) and deleting them in a random sequence in increments of two percent. By eliminating up to 20 percent of members, we attempted to mimic the varying degree of poaching stress likely imposed on wild populations (89). In addition, we were motivated by evidence that many synthetic, biological systems (90) are organized around several, highly connected nodes, important for network development and stability (91). We compared the effect of targeted deletions against a null model by also deleting 20 percent of network members randomly (together referred to as ‘*deletion types*’) in increments of two percent (collectively referred to as ‘*deletion proportions*’). Each deletion proportion was replicated 1000 times per both deletion types and both metrics (92).

After each deletion proportion, in each deletion type and metric, we quantified four, established, theoretical indices diagnostic of social network connectivity and efficiency of social diffusion. These indices included the clustering coefficient and weighted forms of the diameter, global efficiency and modularity. Weighted variants of these indices are informative when individuals associate differently with different conspecifics, which has been reported in elephants (e.g., young adults may associate more frequently with close rather than distant kin) (63). Given the importance of fission-fusion dynamics in elephant populations occurring through interactions among immediate and distant kin (93), we quantified the clustering coefficient and weighted modularity before and after removal of socially influential elephants. By characterizing the number and weight of links within (i.e., clustering coefficient) and across (i.e., modularity) disparate subgroups or modules, we simultaneously comprared the change to network connectivity at the social unit and population levels. By measuring weighted diameter and global efficiency we aimed to illustrate the potential rapidness (i.e., diameter) and pervasiveness (i.e., global efficiency) of social diffusion. Evaluating these indices in the context of elephant social networks allowed us to identify social interactions with capacity for timely and pervasive diffusion of social currency, and their change after poaching-like disturbance. The definitions of these indices and our predictions regarding their change after deletions are detailed in Table 1 (48).

We assessed the mean value of each index as a function of the proportion, type and metric of deletion. Each deletion condition (e.g., targeted deletion of two percent of the most mature network members) was repeated 1000 times — a process theoretically unlimited in the sample size. Therefore, instead of using a comparison of means statistical test informed by a biological distribution, we quantified the difference in the effect size between means of targeted and random deletions using Hedge’s g test (94). We expressed the differences in the mean values between all corresponding conditions using the 95 percent confidence intervals.

### Virtual data

#### Characterizing composition and association properties in virtual populations

To evaluate the impact of poaching-like disturbance on global network structure in the context of sociodemographic diversity likely seen in wild elephant communities, we generated 100 virtual populations based on empirical population composition (78). Each virtual population consisted of females in the previously detailed age categories (Table 1) and four social tiers, namely core, bond, clan and non-kin clan group (S1 Table) (57).

Evaluating the distribution of AIs in the empirically based network, according to age category and kinship, revealed the following patterns. 1) Individuals of any age category were most likely to associate within their core group. They were also more likely to associate with kin from the same bond group than from other bond groups; then with individuals from their clan; and lastly with non-kin (104). 2) In a core group, individuals of any age category were slightly more likely to associate with conspecifics from older age categories (Figure 1a). Since these patterns are generally consistent with the dynamics described in many elephant populations (genetic relatedness — (104, 105); multilevel structure — (78); spatial proximity — (63, 106)), we used the empirically based AI ranges for social network assembly in the virtual populations. To show the parallels, we present the ranges of dyadic associations across all age categories and social tiers in the empirically based and virtual populations (Figure 1 and S1 Table).

**Figure 1.**
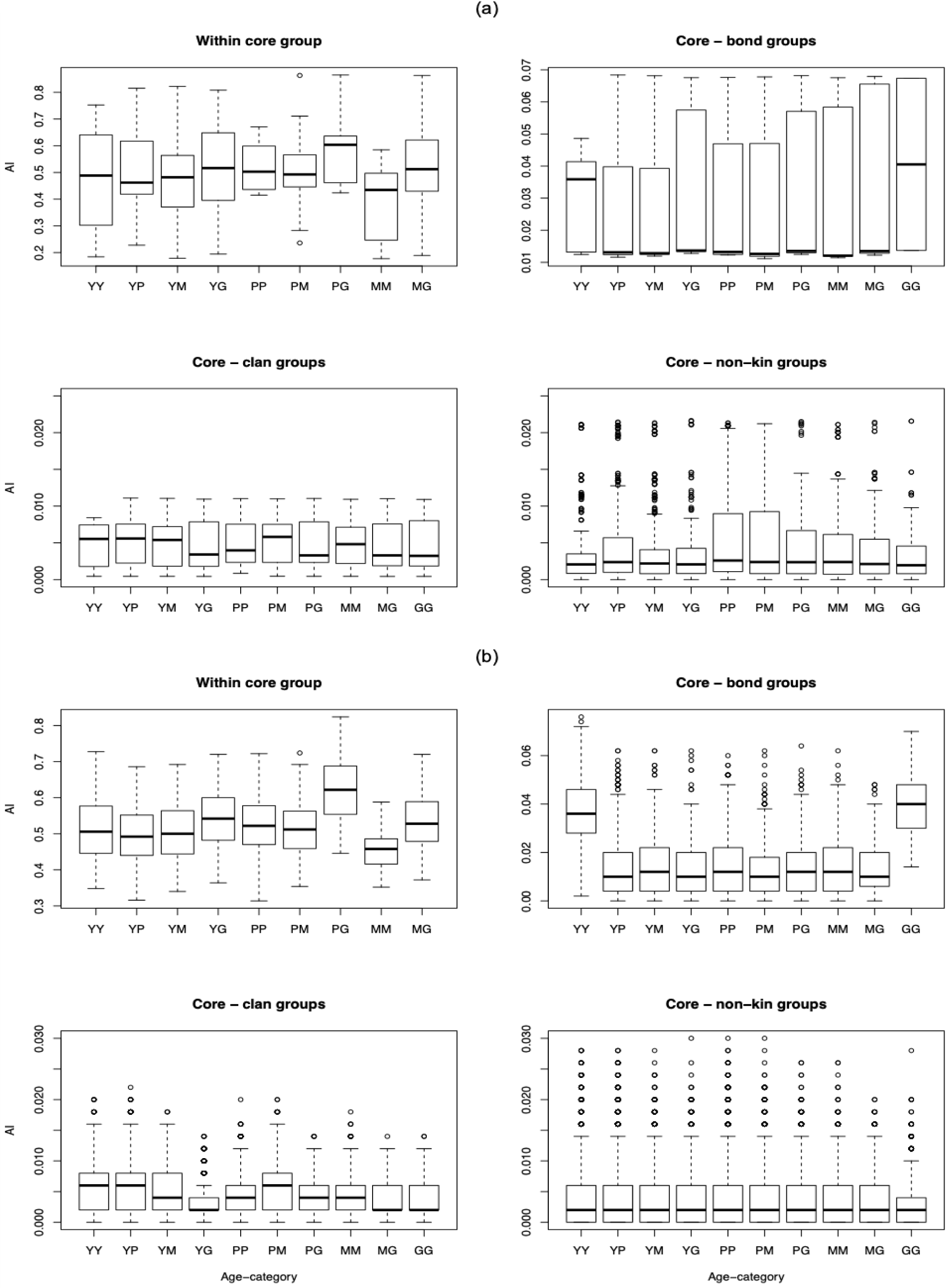
Graph representing the distribution of association indices (AI) in **(a)** the empirically based versus (**b)** virtual populations as a function of age category (Y = young adult; P = prime adult; M = mature adult; G = matriarch) and kinship of the associating individuals. A detailed account of population composition in the empirically based versus virtual populations can be found **S1 Table** in the Supplemental Materials section.

#### Simulating virtual social networks

To simulate 100 virtual social networks, we used a spatiotemporally nonexplicit, individual-based model at two levels — between core groups and then dyads. The range of probabilities of kinship- or age-based association between two groups or individuals, respectively, were drawn from a triangular distribution mimicking empirically based data (Figure 1b). At each time step, each dad in the population had the opportunity to associate. Once a core group and a dyadic association had been determined to occur, the time step was terminated and the total number of observed associations per each dyad was updated (S1 Figure).

The networks had started to reach a plateau after 500-time steps (S2 Figure). However, to study how deletions may affect the global structure of networks at different stages of development, we stopped the simulation at 100-, 200-, 300-, 400- or 500-’*time steps’*. From these networks, we noted the age category and quantified betweenness of every individual. To compare their structure, we present graphs of the empirically based network and an example of a similarly sized virtual network (Figures 2). They appear similar in age category makeup and WGS associations. The empirically based network has fewer BCG associations and nodes with higher overall betweenness values than the virtual network.

**Figure 2.**
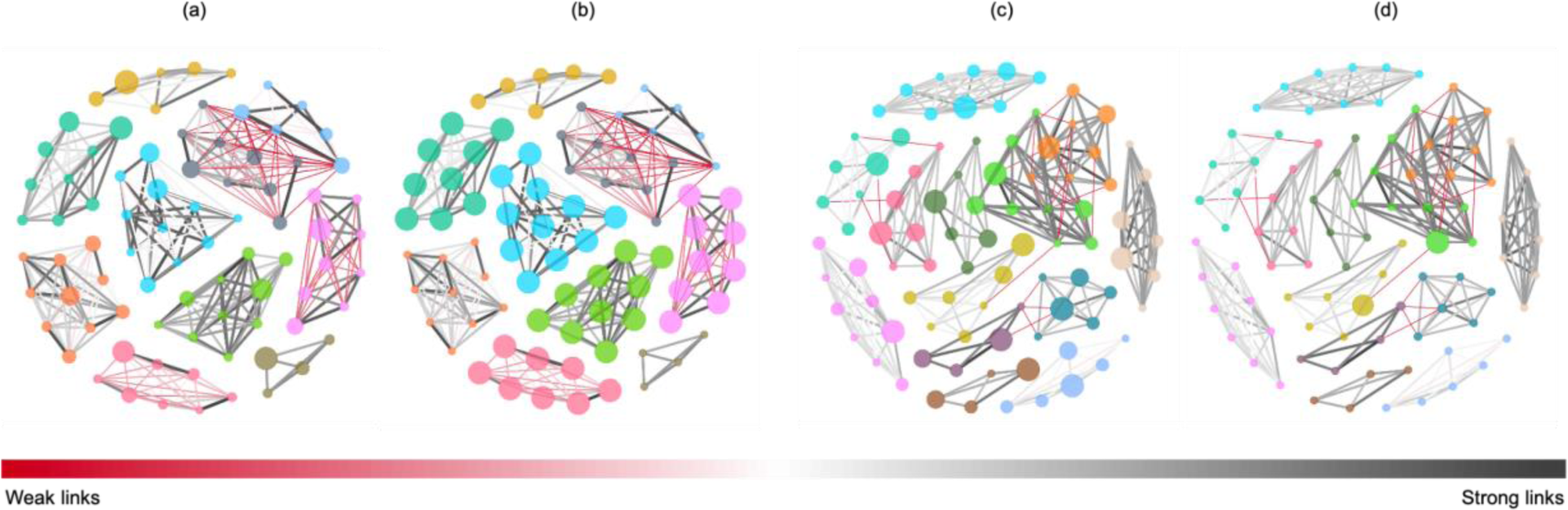
Social network graphs of the empirically based population with color partitioning according to a core group, considered from the perspective of either **(a)** age category or **(b)** betweenness; and a comparable example of a virtual population with the partitioning according to a core group, and either **(c)** age category or **(d)** betweenness. The nodes are ranked by size where the largest nodes indicate oldest age or highest betweenness. The links are ranked according to their relative weight. The color and thickness scheme depicting the weight of each link ranges from red/thin (low) to to dark grey/thick (high weight). The links with weight less than 5 percent were filtered out for visual clarity.

#### Conducting deletions using virtual social networks

To measure if the disappearance of the most socially influential individuals changed the connectivity and efficiency in the 100 virtual networks at each of the five time steps, we performed a series of targeted and random deletions. Individuals were deleted in four percent increments, ranging from zero to 20 percent. In targeted deletions, 20 percent of individuals selected for removal had the highest betweenness or belonged to the oldest age category. During each random deletion, the same proportion of individuals as in targeted deletions was removed randomly, disregarding their betweenness or their age category. After every deletion proportion, we recalculated the following network level indices: clustering coefficient, as well as weighted diameter, global efficiency and modularity (Table 1). As in the empirically based portion of our study, we used the Hedge’s g test to quantify the difference in the effect size between the means of all network indices across 1) the deletion proportion spectrum, 2) deletion type, 3) time step and 4) deletion metric (94).

Motivated by a preliminary assessment indicating a high degree of resilience to fragmentation after the deletion of the oldest or most central members, even at early stages of network formation (i.e., 100-time steps), we explored if simulated networks would break down when subject to prior elimination of relatively weak associations (107). Here we wanted to determine if weak associations, likely formed among individuals with high betweenness, could also be explained by age category. During this process, we manipulated only the most robust networks (i.e., 500-time steps) by filtering out the ‘*weakest links*.’ To do so, we divided the value of each link in the association matrix by the highest link value and eliminated the links with values up to three percent of the highest link in increments of one percent. After each elimination without replacement, we carried out the deletions and quantification of outcomes as described above.

The social network quantification and analysis of both the empirically based and virtual data were performed using the R statistical software, version 3.2. (R Core Team 2017). Visualization of the social networks was performed in Gephi software, version 0.9.2 (108).

## Results

### Empirically based network

Contrary to our expectations, the results of targeted deletions in the empirically based portion of our study revealed disparities in almost all network indices between age category and betweenness (S2 Table) and an overall unexpected level of resilience against disturbance.

The effect size statistics estimating the mean difference between age category-targeted and random deletions at each deletion proportion revealed no change in clustering coefficient, as well as weighted global efficiency and modularity. Weighted diameter decreased in targeted deletions but only at larger deletion proportions (e.g., proportions in the interval {0.1, 0.2}) (Figure 3). Although we did not expect these results, the removal of the oldest elephants in simulated populations appears less damaging to the network connectivity than we expected. Network efficiency, however, based on the weighted diameter results, was negatively affected by elimination of seniors.

**Figure 3.**
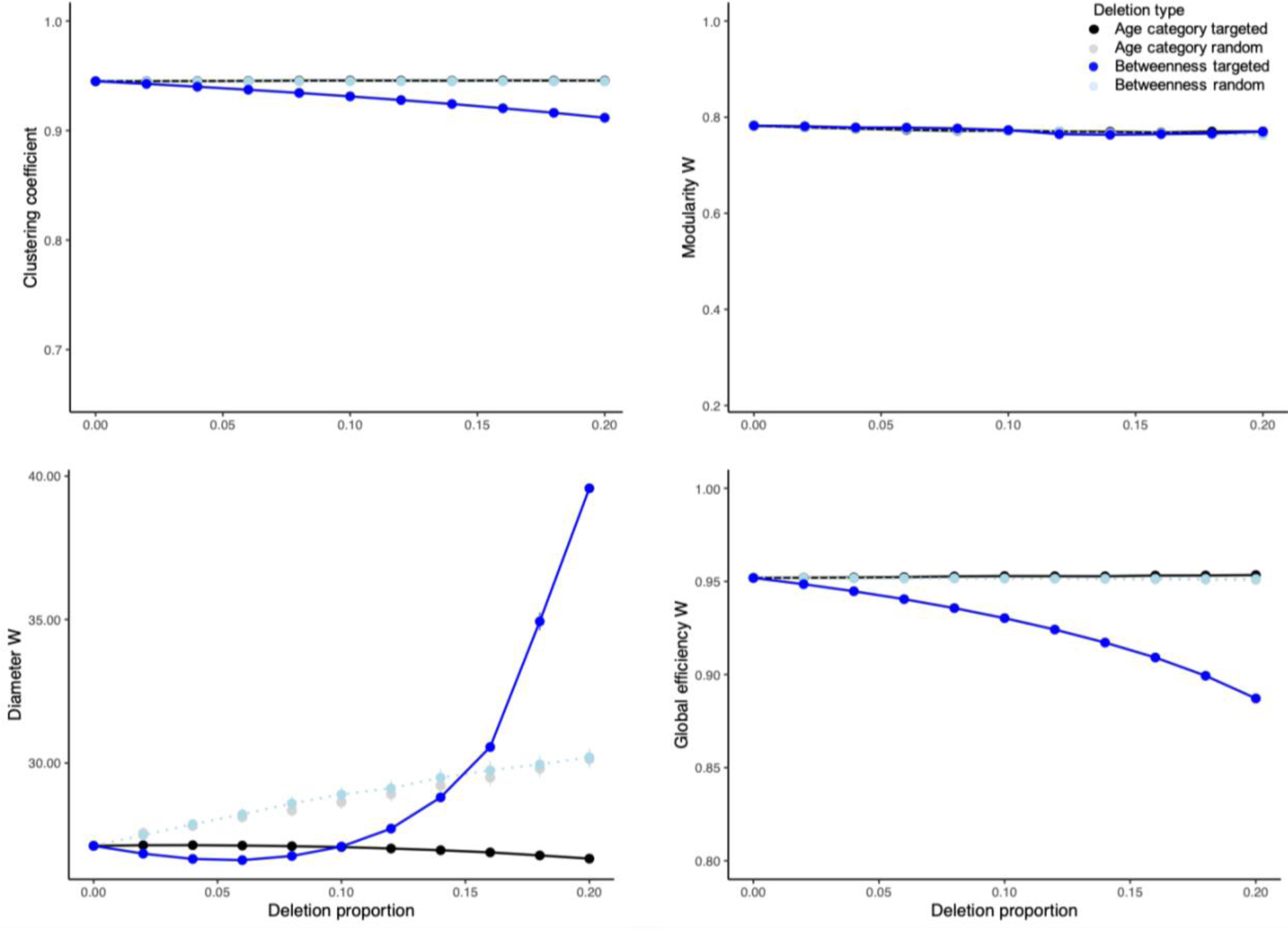
Graphs representing results (mean plus 95% confidence interval) of 1000 deletions per each combination of deletion proportion (i.e., 0-20%) and type (i.e., random vs. targeted) in the empirically based network. The deletions were either targeted according to age category (black series) or betweenness (blue series); or were random (grey and teal series represent random deletions without considering individual traits conducted as control conditions to age- or betweenness-targeted experiments, respectively). The network indices evaluated included clustering coefficient as well as weighted modularity, diameter and global efficiency. For across-species context, the minima of y-axis ranges per clustering coefficient as well as weighted modularity and global efficiency are plotted to express the minima from a similar, theoretical treatment in an egalitarian primate society (92). The weighted diameter index depends on group size, thus the pertinent y-axis is not expressed in across-species context. For results of Hedge’s g test expressing the difference in the effect size between the mean values of each network level index in targeted versus random deletions along the deletion proportion axis and per deletion type, refer to **S2 Table** in the Supplemental Materials section.

In contrast, the effect size statistics comparing the differences between targeted and random elimination of individuals with highest betweenness, as a function of deletion proportion, showed an expected decrease in clustering coefficient and weighted global efficiency, as well as an increase in weighted diameter (Figure 3). Weighted modularity revealed no change relative to random deletions (Figure 3). This set of results indicates that the loss of the most central conspecifics, particularly if more than 10 percent of them are removed, impedes connectivity and efficiency in simulated networks.

### Virtual networks

The results in the virtual portion of this study were similar to those from the empirically based portion. When age category was the focus of deletions, the effect size statistics comparing means of targeted and random deletions in the 100 virtual networks, along the time step and deletion proportion axes, revealed an increase in clustering coefficient and weighted global efficiency, and a decrease in weighted diameter. For the latter two indices, large effect size statistics were only apparent at early time steps and large deletion proportions (e.g., up to 300- time steps and proportions in the interval {0.16, 0.2}). There was no change in mean, weighted modularity between targeted and random deletions (S3 Table). Contrary to our expectation, these results suggest that removal of older individuals improved connectivity in 400- and 500- time step networks but without improving their efficiency.

When targeted deletions were performed according to betweenness, the clustering coefficient and weighted global efficiency decreased, while weighted modularity and diameter increased. The effect size statistics for these indices were large across most time steps and deletion proportions (S3 Table). As we expected, these results point to a decrease in connectivity and efficiency of simulated networks and importance of individuals with high betweenness in shaping these network features.

Elimination of the weakest association links with values ranging from one to three percent of the highest link in 500-time step networks led to multiple events of breakdown into at least two modules (S4 Table). Given their ‘premature’ disruption, we excluded these networks from the subsequent deletions. In the remaining filtered networks, targeted deletions of individuals with the highest betweenness, more so than age category, caused more fragmentation than random deletions. Finally, although the weakest links were rather evenly distributed between individuals of various age categories, they occurred more often among individuals from different clans (S3 Figure) indicating an important role in network connectivity.

## Discussion

In this study, we addressed a timely question about the response of animal groups to human disturbance by simulating poaching in African savanna elephant populations. After targeted removal of socially influential individuals, according to their age category or position in a social network (i.e., betweenness), we characterized network indices associated with cohesion and transfer of information in animal groups. We anticipated that targeted disturbance would 1) perturb theoretical indices of network connectivity and the efficiency of social diffusion immediately after disturbance and 2) increase as a function of deletion proportion (i.e., 0 - 0.2) leading to network breakdown.

Contrary to our expectations, targeted deletions according to age category resulted in improved connectivity in simulated networks. This outcome, however, instead of pointing to social influence of seniors, revealed their peripheral roles in contributing to network connectivity relative to younger conspecifics. Elimination of individuals with high betweenness led to an anticipated decrease in indices expressing connectivity and efficiency of social diffusion in simulated networks. Unlike age category, betweenness proved to be an indicator of social influence in the context of strong links among close kin as well as weak links among distant kin. Finally, regardless of the deletion metric, the simulated networks did not break down even when subject to relatively high degree of ‘poaching’, leaving the question of a theoretical breaking point outstanding.

The disparities between age category- and betweenness-specific deletions are consistent with intraspecific behaviors in species with multilevel sociality, established dominance hierarchy and high degree of tolerance towards subordinate group members (109). For instance, in real elephant populations, immature individuals are rather indiscriminate in their affiliations and likely to engage with multiple conspecifics of different ages and kinship (58,59,110). Frequent bouts of social engagement may afford them some social skills without direct engagement of senior kin and fosters cohesion between distinct subgroups (31, 73). In contrast, similarly to mature individuals in other group-living species (111, 112), senior elephants may be more selective about their social partners and less sociable (78). Their value as social intermediaries contributing to network connectivity and efficiency may for that reason be comparable to their immature conspecifics (36, 73), regardless of the wealth of socioecological experience seniors likely possess and display during social activities (e.g., such as group antipredator defense led by the matriarch — (113)).

This type of organization, where network stability is mediated by different categories of individuals, exemplifies a decentralized system, likely selected for to buffer destabilizing effects of prolonged fission or stochastic events such as disease-induced die-off (114) or poaching. The notion of network decentralization, reflected in our simulation, parallels the findings by Goldenberg and collaborators who propose that the redundancy between social roles of mature elephants, prior to poaching, and their surviving offspring is a potential mechanism of network resilience against breakdown (73). The simulated networks in our research were also resilient to removal of the socially influential group members. Given the seemingly greater flexibility and interconnectedness in elephant populations, relative to other closely knit social species (92) finding hypothetical limitations to social resilience may require evaluating more intensive yet biologically meaningful ‘poaching’ disturbance than considered in our work (115).

Although our assessment of the effects of disturbance on social organization and resilience does not account for the dynamic or indirect responses to poaching (e.g., network reorganization or avoidance of poaching hotspots), it is a valuable first step in systems with limited real-world data. Having access to information about the proportion and type of missing group members may 1) offer basic but meaningful insights about why some poached elephant populations take exceptionally long to recover from member loss (116), while others recover much quicker(117) and 2) help reason about the fate of recovering populations. Our ideas may also be transferable to management of other group-living, keystone species (118–122). For instance, applied without consideration for social interactions, trophy hunting of pride lions may intensify infanticide by immigrant males (23,28,120) and displace distressed females to hunt in fringe habitats exacerbating conflict with humans (121, 123). Prior to making decisions about lethal management or translocations of ‘problem’ individuals, wildlife managers may be well served by simulating relevant disturbance on focal populations, quantifying social network effects and adjusting management decisions for better outcomes (39, 124). As another example, the use of ASNA in captive animal populations is already helping researchers characterize the dynamics of harmful agonistic interactions, such as tail biting in newly mixed groups of domestic pigs (125). These data may help parametrize simulated disturbance to social network structure in captive systems by taking into account traits such as genetic relatedness in group composition to determine its link to aggression and health of animal subjects. Insights from this type of assessment may improve animal husbandry and safety of farm workers (126, 127).

In summary, our work confirms previous findings that although elimination of the most central network members decreases network connectivity at the population level, it does not lead to network fragmentation. Uniquely, however, our research shows that poaching-like stress in a large number of virtual elephant populations impedes the theoretical efficiency of social diffusion. A follow-up question about the relationship between the structural network changes and population performance will require simulating a dynamic process that accounts for network reorganization after poaching. In addition, to tease apart an individual’s importance due to network position versus age-specific experience will require a method that accounts for interaction-mediated information transfer. Still, our simulation platform can be easily altered to test basic hypotheses about disturbance of social interactions in wild and captive systems.

## Data Availability Statement

Data and code will be available on the Dryad repository.

## Acknowledgments

Funding for this project was provided by the 2018-2019 STEM Chateaubriand Fellowship Program. Empirical data were provided by the Amboseli Trust for Elephants.

## Supporting Information Captions

**S1 Table.** The summary composition of 100 virtual populations with the numbers of clan, bond and core groups, as well as individuals per population; the number of bond and core groups, and individuals per clan; the number of core groups per bond group; and the number of individuals per bond and core groups. The distribution of age categories within each core group was the following: young adults (mean = 2 individuals, min = 1, max = 5); prime adults (mean = 2, min = 0, max = 7); mature adults (mean = 1, min = 0, max = 3); and matriarchs (mean = 1, min = 1, max = 1). The composition of the empirical population is included as a reference (i.e., = 10 core groups including a total of n= 83 individuals) (78, 88).

| Demographic group | Minimum | Maximum | Median | Empirical contrast |
| --- | --- | --- | --- | --- |
| Clan groups per population | 1 | 8 | 5 | 3 |
| Bond groups per population | 1 | 28 | 14 | 8 |
| Core groups per population | 5 | 86 | 40 | 10 |
| Bond groups per clan group | 1 | 5 | 3 | 4,3,1 |
| Core groups per clan group | 1 | 20 | 9 | 5,4,1 |
| Core groups per bond group | 1 | 5 | 3 | 1,1,2,1,2,1,1,1 |
| Individuals per population | 95 | 760 | 350 | 83 |
| Individuals per clan group | 10 | 175 | 74 | 39,36,8 |
| Individuals per bond group | 1 | 45 | 25 | 10,6,19,4,16,11,9,8 |
| Individuals per core group | 4 | 15 | 8 | 10,6,9,10,4,6,10,11,9,8 |

**S2 Table.** Results of Hedge’s g test expressing the effect size difference between mean values of each network index in targeted versus random deletions in empirically based networks, along the deletion proportion axis, with deletions performed according to either age category or betweenness (94). Bold values indicate medium (≥ |0.5|) and large (≥ |0.8|) effect size.

| Network Index | Deletion proportion | Hedge's g statistic |  |
| --- | --- | --- | --- |
|  |  | Age category | Betweenness |
| Modularity W | 0.02 | -0.0348 | 0.2513 |
|  | 0.04 | -0.0239 | 0.1394 |
|  | 0.06 | -0.1002 | 0.3639 |
|  | 0.08 | -0.0538 | 0.2219 |
|  | 0.1 | 0.0380 | 0.1154 |
|  | 0.12 | 0.0171 | -0.4311 |
|  | 0.14 | 0.0630 | -0.1442 |
|  | 0.16 | 0.0315 | -0.1178 |
|  | 0.18 | 0.2038 | 0.0303 |
|  | 0.2 | 0.3683 | 0.3449 |
| Global efficiency W | 0.02 | 0.0750 | **-1.5411** |
|  | 0.04 | 0.1247 | **-2.2205** |
|  | 0.06 | 0.1565 | **-2.9173** |
|  | 0.08 | 0.2054 | **-3.5236** |
|  | 0.1 | 0.2066 | **-4.04176** |
|  | 0.12 | 0.1941 | **-4.6401** |
|  | 0.14 | 0.1994 | **-5.3114** |
|  | 0.16 | 0.2650 | **-6.0381** |
|  | 0.18 | 0.2883 | **-6.9214** |
|  | 0.2 | 0.3328 | **-8.1713** |
| Clustering coefficient | 0.02 | 0.0476 | **-1.6673** |
|  | 0.04 | 0.0904 | **-2.3356** |
|  | 0.06 | 0.1218 | **-3.0060** |
|  | 0.08 | 0.1693 | **-3.5128** |
|  | 0.1 | 0.1572 | **-3.9375** |
|  | 0.12 | 0.12531 | **-4.3778** |
|  | 0.14 | 0.1212 | **-4.8515** |
|  | 0.16 | 0.1635 | **-5.3056** |
|  | 0.18 | 0.1570 | **-5.7977** |
|  | 0.2 | 0.1709 | **-6.2864** |
| Diameter W | 0.02 | -0.2706 | -0.4453 |
|  | 0.04 | -0.3439 | **-0.5870** |
|  | 0.06 | -0.4264 | **-0.6470** |
|  | 0.08 | -0.4898 | **-0.6503** |
|  | 0.1 | **-0.5604** | **-0.5966** |
|  | 0.12 | **-0.6311** | -0.4333 |
|  | 0.14 | **-0.7000** | -0.1889 |
|  | 0.16 | **-0.7693** | 0.1999 |
|  | 0.18 | **-0.8560** | **0.9766** |
|  | 0.2 | **-0.9446** | **2.4932** |

**S3 Table:**
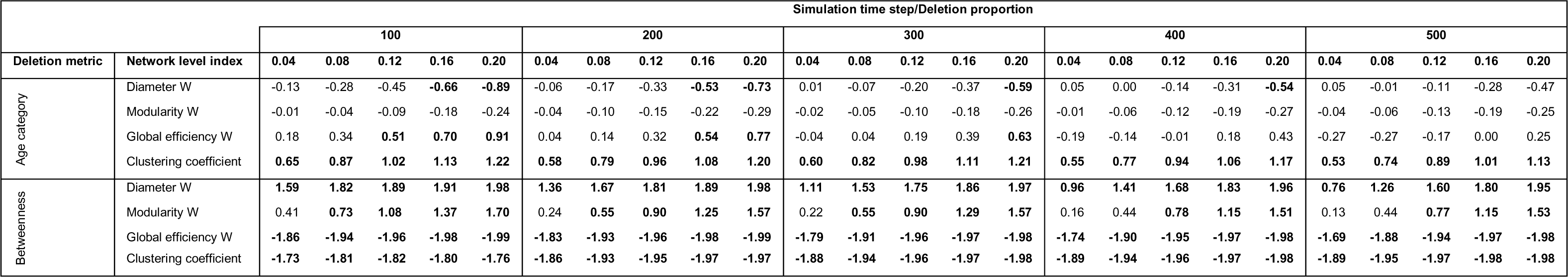
Results of Hedge’s g test expressing the difference in the effect size between the means of each network level index (i.e., weighted (W) diameter, modularity and global efficiency; as well as clustering coefficient) in targeted and random deletions, spanning all network time step and deletion proportion increments. The deletions were performed according to age category or betweenness (94). Bold values indicate medium (≥ |0.5|) and large (≥ |0.8|) effect size.

**S4 Table.** The summary of the percentages of filtered, virtual networks that broke down into two or more modules as a result of the deletions performed according to age category or betweenness. The filtering process was carried out before the onset of the deletions by dividing the value of each link in the association matrix by the highest link value and eliminating the links with values up to three percent of the highest link in increments of one percent (107). Only 500-time step networks were considered in these experiments.

| Deletion metric | Deletion type | Filtering percent | Deletion proportion |  |  |  |  | Minimum number of modules at 0.2 deletion | Maximum number of modules at 0.2 deletion |
| --- | --- | --- | --- | --- | --- | --- | --- | --- | --- |
|  |  |  | 0.04 | 0.08 | 0.12 | 0.16 | 0.2 |  |  |
| Age category | Targeted | 1 | 0 | 0 | 0 | 0 | 0 | 1 | 1 |
|  |  | 2 | 0 | 1 | 2 | 2 | 2 | 1 | 1.41 |
|  |  | 3 | 0 | 0 | 0 | 0 | 0 | 1 | 1 |
|  | Random | 1 | 0 | 0 | 0 | 0 | 0 | 1 | 1 |
|  |  | 2 | 3 | 4 | 5 | 8 | 14 | 1 | 1.22 |
|  |  | 3 | 100 | 100 | 100 | 100 | 100 | 1.25 | 1.25 |
| Betweenness | Targeted | 1 | 0 | 0 | 0 | 0 | 0 | 1 | 1 |
|  |  | 2 | 6 | 14 | 17 | 19 | 19 | 1 | 4 |
|  |  | 3 | 100 | 100 | 100 | 100 | 100 | 2 | 5 |
|  | Random | 1 | 0 | 0 | 0 | 0 | 0 | 1 | 1 |
|  |  | 2 | 1 | 5 | 7 | 11 | 16 | 1 | 1.34 |
|  |  | 3 | 0 | 100 | 100 | 100 | 100 | 1.22 | 1.22 |

**S1 Figure.**
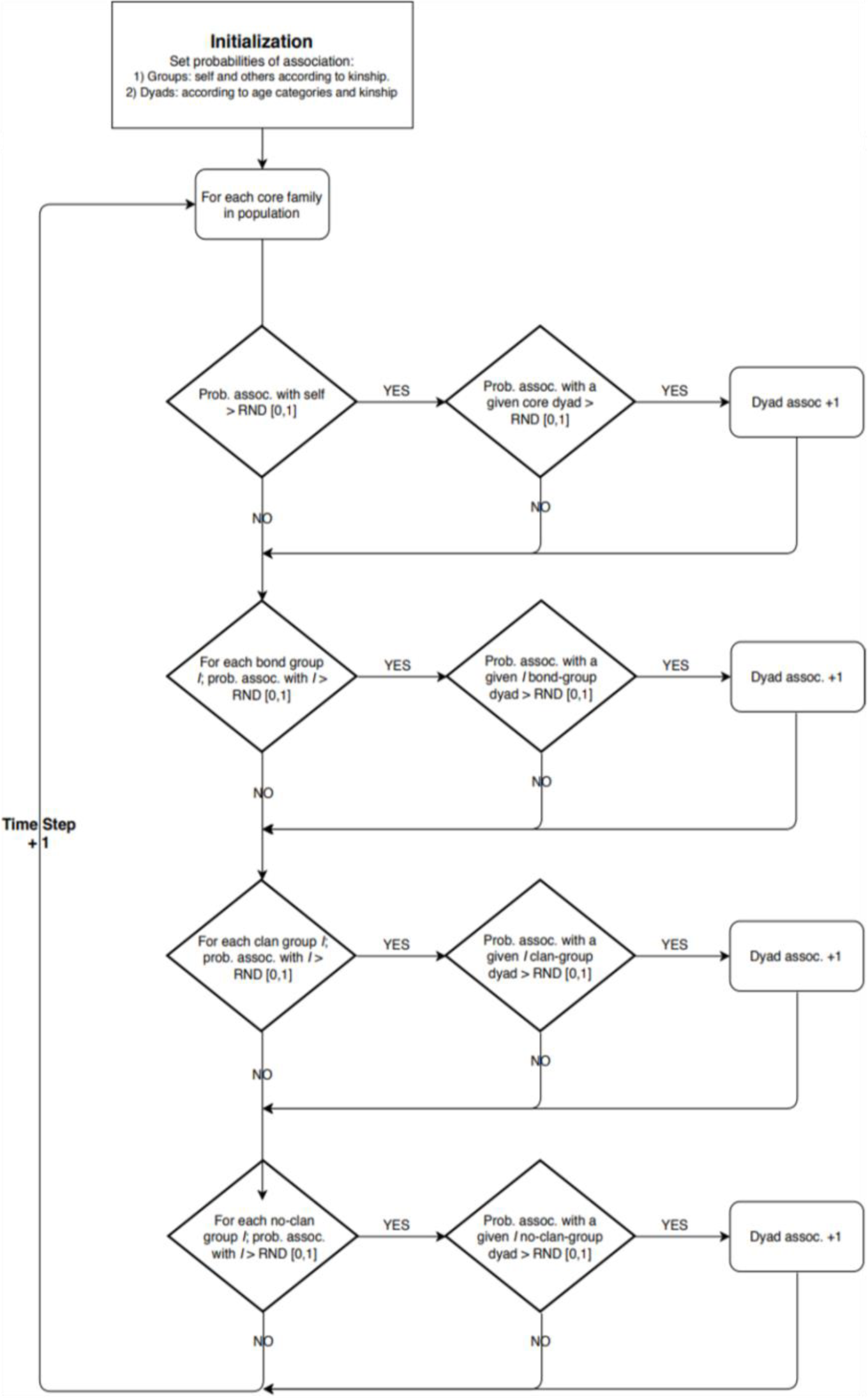
Flow chart summarizing the process of simulating social networks among virtual elephant populations. At initialization, the probabilities of association between and within groups are set according to kinship and age category (Figure 1). At the beginning of each time step, the set probability of association between or within each set of groups and between each dyad is compared to a randomly generated number (RDN) between {0,1}. If this probability is greater than RDN, the association is set to occur; if this probability is lower than RDN, the association does not occur, and the time step is terminated. At the end of each time step the number of times a specific dyad has formed across all previous time steps is updated (i.e., increased by one if the association had occurred, or remained the same otherwise). For the distribution of network indices as a function of the number of simulation time steps refer to **S2 Figures**.

**S2 Figure.**
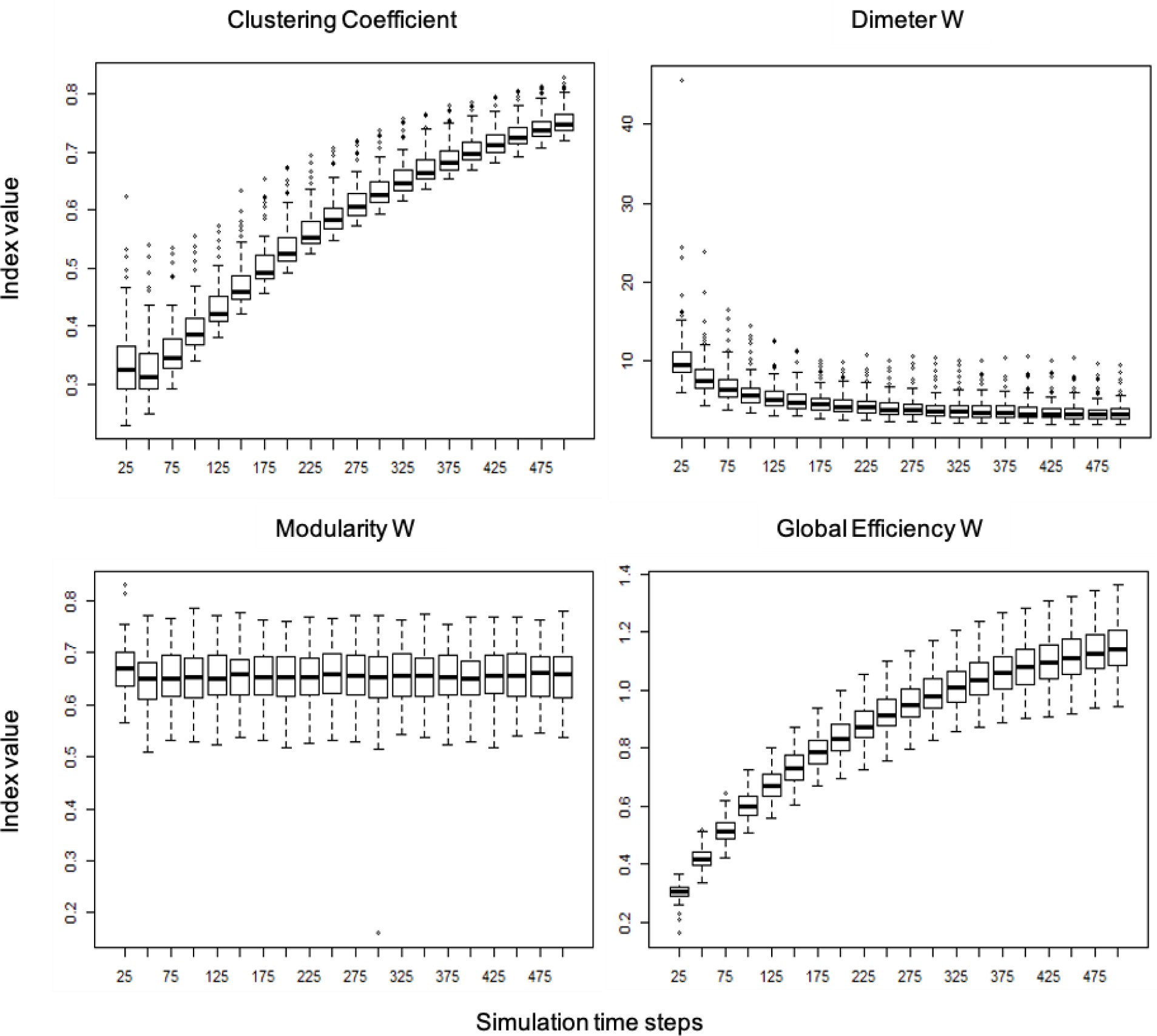
The distribution of values per each of the network indices evaluated, including the clustering coefficient, as well as weighted diameter, global efficiency and modularity, expressed as a function of the number of simulation time steps. The 500-time step cut-off was based on when the density (or the proportion of existing interactions among network members, relative to the number of possible interactions) of the resulting networks started to reach a plateau (∼ 75% median density) (80).

**S3 Figure.**
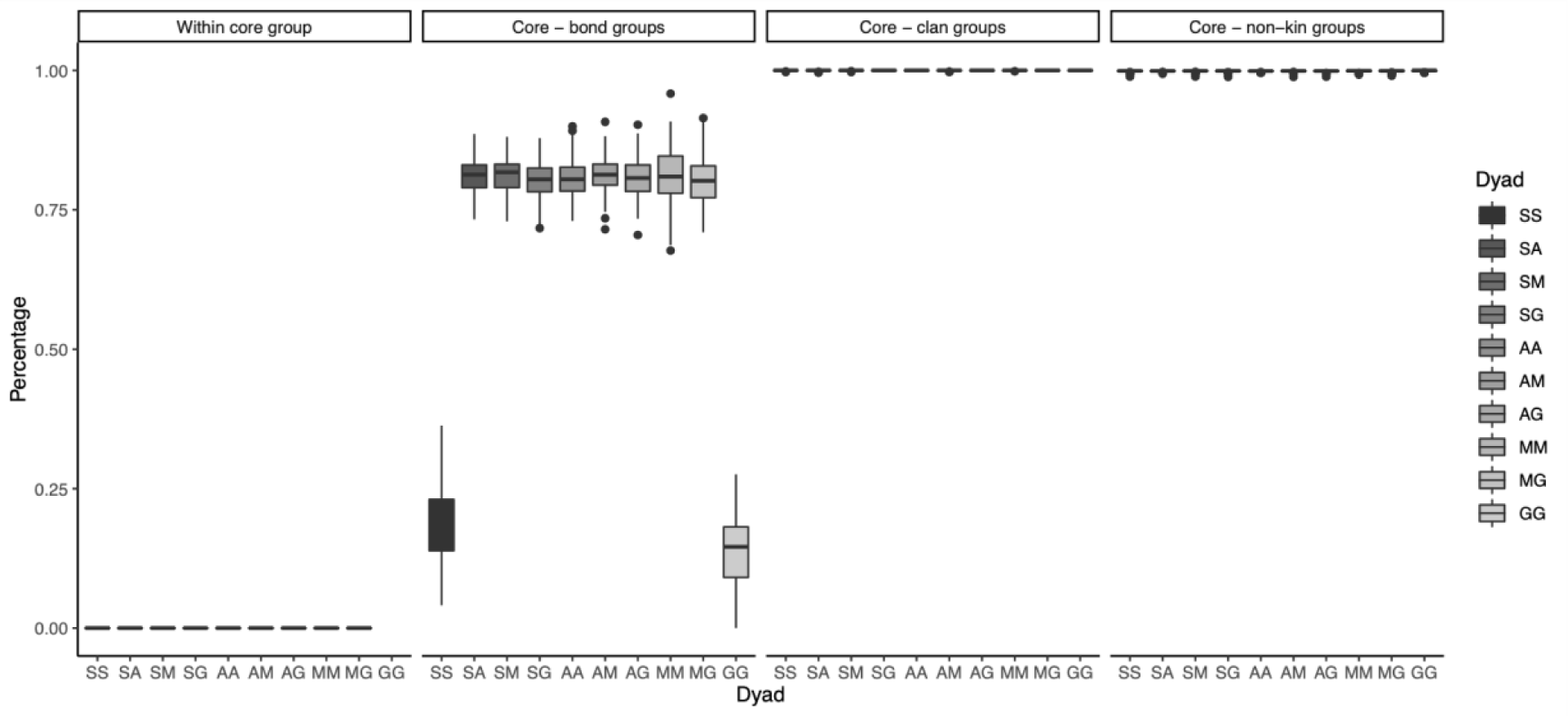
The percentage of the weakest associations (i.e., links with values up to three percent of the highest link) filtered out from the 500-time step, virtual networks prior to deletion experiments. These links are presented according to age class in a dyad (Y = young adult; P = prime adult; M = mature adult; G = matriarch) and one of four social tiers. For the summary of filtering experiments showing percentages of filtered, 500-time step, virtual networks that broke down into two or more modules as a result of the deletions performed according to age category or betweenness, refer to **S4 Table**.

